# Gene-centric analysis of *Raskinella chloraquaticus* reveals a functionally conserved taxonomic group in global drinking water distribution systems

**DOI:** 10.1101/2025.11.10.687676

**Authors:** Ashwin S Sudarshan, Konstantinos T Konstantinidis, Ameet J Pinto

**Affiliations:** School of Civil and Environmental Engineering, Georgia Institute of Technology, Atlanta, GA 30332, USA; School of Earth and Atmospheric Sciences, Georgia Institute of Technology, Atlanta, GA 30332, USA

**Keywords:** Drinking water microbiology, Metagenomics, Pangenomics, Recombination

## Abstract

A recent metagenomic survey of drinking water systems revealed that a highly prevalent and dominant uncultured bacterial genus (*Raskinella*) was represented globally by a single species (*Raskinella chloraquaticus*). *R. chloraquaticus* comprises of two sub-species groups, Lineages 1 and 2, the former representing a globally prevalent genomovar. The objective of this study was to perform comparative analysis of the gene content of *R. chloraquaticus* to characterize the gene-level diversity and determine factors shaping the diversity of this species. Pangenome analysis revealed that *R. chloraquaticus* possesses a core set of genes that constitute a major portion (87.74%) of the known gene content of the genome. Furthermore, most of the gene diversity of *R. chloraquaticus* is associated with Lineage 2 organisms, which consists of at least four distinct genomovars. Lineage 1 organisms consist of a higher proportion of identical genes than would have been expected if changes primarily occurred through random mutations and thus is potentially indicative of recombination. In contrast, Lineage 2 organisms appear to have emerged through random mutations and display stronger geographic preference. These results indicate that homologous recombination and geographic isolation likely shape the genetic repertoire of *R. chloraquaticus*. Further, the high level of gene conservation in *R. chloraquaticus* may be reflective of highly selective environment in drinking water systems. Thus, *R. chloraquaticus* may represent a model organism to probe selective pressures shaping the drinking water microbiome.

## Introduction

Drinking water systems harbor diverse and complex microbial communities despite multibarrier treatment to reduce microbial concentrations (Proctor and Hammes 2015). Previous research has indicated that the persistence of the drinking water microbiome can be attributed to a variety of physiological and ecological traits ranging from enhanced disinfection resistance to ecological sheltering in biofilms (Prest et al. 2016). Recently, we constructed a comprehensive drinking water genome catalogue to broadly evaluate the biogeography of the drinking water microbiome and identify core genera that were prevalent across drinking water distribution systems (DWDS) globally. Through this analysis, we identified an uncultured genus (*Raskinella*) that was prevalent and globally dominant within disinfected DWDS (Sudarshan et al. 2024).

*Raskinella* was unique compared to other well-known drinking water-relevant genera (e.g., *Sphingomonas*, *Mycobacterium*, *Hyphomicrobium*, etc.) in that a single species (*Raskinella chloraquaticus*) within this genus was observed in all systems where this genus was detected. This was in contrast to other frequently detected genera like *Sphingomonas* and *Mycobacterium* where 34 and 25 different species within these genera were detected, respectively. Within *R. chloraquaticus*, we observed two distinct sub-species groups (henceforth, referred to as lineages) based on average nucleotide identity (ANI) differences, which were referred to as Lineage 1 and Lineage 2. Lineage 1 represented a globally prevalent genomovar (ANI >99.5%) and was the more prevalent *R. chloraquaticus* lineage. In contrast, Lineage 2 was a more diverse lineage group (ANI range: 96.7-99.91%) and was less prevalent than Lineage 1. Metabolic annotation and modeling indicated that *R. chloraquaticus* may grow necrotrophically, synthesize homoserine lactones, and has the ability to degrade halogenated compounds. The ability to utilize decay products from inactivated microorganisms, form biofilms, and utilize halogenated organics may impart a distinct ecological advantage to *R. chloraquaticus* in the context of disinfected drinking water systems. Metabolic analysis also revealed key differences between the two lineages of *R. chloraquaticus* which could explain differences in their prevalence. For example, Lineage 1 possessed the ability to oxidize thiosulfates which was not observed in Lineage 2. The extremely low genetic diversity within *Raskinella* and the global prevalence of *R. chloraquaticus* in drinking water systems necessitates an investigation into the genomic factors that may underpin its persistence and global distribution.

Genome-centric analysis using assembled metagenome assembled genomes (MAGs) can not only provide phylogenetic placement but also comparative insights into their functional potential. The latter can be achieved by performing a comparative analysis using multiple independently assembled MAGs within the environmental context from where they were assembled. This analysis can not only help determine gene-level diversity of closely related MAGs but also shed light on the importance of these genes towards MAG prevalence. Furthermore, gene-centric analyses can also capture potential functional diversity across different MAGs which is not captured through whole-genome comparisons which rely on sequence identity. High gene diversity could indicate an ability to thrive in diverse habitats while lower diversity would suggest adaptation to a relatively narrow ecological niche (e.g., disinfected drinking water systems). Therefore, characterizing the gene-level diversity can shed light on the broad traits of *R. chloraquaticus* and the genes underpinning these traits (Medini et al. 2005). Furthermore, demarcating the genetic boundaries between closely related organisms is necessary to delineate the factors that contribute to the maintenance of species groups within complex microbial communities. Previous studies have suggested both molecular factors (homologous recombination, diversifying mutation) and ecological or extrinsic factors (ecosystem, geographic isolation) can play a role. Such insights are particularly important for organisms like *R. chloraquaticus*, considering it exhibits minimal genomic variability (ANI > 99.5%) in disinfected drinking water systems globally.

Therefore, the overall goal of this study was to perform a comparative analysis of the gene content of *Raskinella chloraquaticus* to answer the following questions: (a) what is the gene-level diversity of the two closely related lineages within *R. chloraquaticus*? and (b) what are the putative factors responsible for shaping and maintaining the population structure within *R. chloraquaticus* in disinfected drinking water systems?

## Materials and Methods

### Source of Raskinella Chloraquaticus MAGs

*Raskinella chloraquaticus* MAGs were obtained from the Drinking Water Genome Catalog (DWGC) (Sudarshan et al. 2024). Briefly, quality filtered reads from 85 DWDS were individually assembled into contigs which were subsequently binned using multiple binning tools to obtain the MAGs. Assembly and binning were performed individually for each DWDS, and MAGs were dereplicated to obtain the final set of representative genomes for the DWGC database. *R. chloraquaticus* MAGs were manually curated based on quality score (Completeness-5*Contamination) and ANI values to identify non-redundant MAGs (ANI<99%) from each DWDS. This resulted in 42 MAGs which were used for subsequent analysis. The completeness and contamination values for these *R. chloraquaticus* MAGs were determined using CheckM2 v0.1.3 (Chklovski et al. 2023). Genome assembly statistics for the MAGs were determined using Seqkit v2.3.0 (Shen et al. 2024). Gene counts and its quality were determined from summary files from prodigal v2.6.3 through Anvi’o v8 (Eren et al. 2021) (**Table S1**).

### *R. chloraquaticus* relative abundance estimation

Reads from selected DWDS were mapped to the representative MAGs from the two lineages of *R. chloraquaticus* (see Results section) using the CoverM v0.7.0 pipeline (Aroney et al. 2025). Briefly, reads were mapped using bwa-mem (Li 2013) and the trimmed mean average depth (TAD90) value and covered fraction for each MAG across the DWDS was determined using mapped reads that had a minimum percent sequence identity of 97% and a minimum read alignment of 75%. The TAD90 values refer to the coverage of the MAG after excluding the top 5% and bottom 5% of coverage values from all reads mapping to the MAG. TAD90 is an effective metric to describe average depth and corrects for inaccuracies associated with mapping outliers emerging from conserved regions and genomic regions inadequately captured through sequencing. The read based average nucleotide identity (ANIr) values for the different DWDS were calculated using custom scripts from the following public repository [https://github.com/rotheconrad/00_in-situ_GeneCoverage] (Lindner et al. 2024). For ANIr calculation, read mapping was performed using Magic Blast as outlined in the workflow and the read identity cutoff used to determine depth and ANIr was 95%.

### *R. chloraquaticus* pangenome analysis

*R. chloraquaticus* MAGs were imported into Anvi’o v8, and their genes were clustered using the mcl-inflation parameter=10, minbit parameter=0.5, excluding partial gene calls and utilizing DIAMOND (Buchfink et al. 2015) to construct a pangenome. The resulting gene clusters were annotated using the COG20 (Tatusov et al. 2000), Pfam v37.2 (Mistry et al. 2021), KEGG, KOfam (Kanehisa et al. 2023) and CAZy v11 (Yin et al. 2012) databases within Anvi’o v8. Core gene clusters were determined using mOTUpan (Buck et al. 2022) using the command anvi-script-compute-bayesian-pan-core using a bootstrap value of 1000. Gene diversity within *R. chloraquaticus* (and its lineages) were calculated using two metrics: Heap’s law parameter and genome fluidity. Heap’s law describes a relationship between the number of unique genes within a taxonomic group and the number of genomes used within the group. Genome fluidity evaluates the number of unique gene families between two genomes as a fraction of the total gene content in the two genomes. Heap’s law parameter and genome fluidity were calculated using a custom script using R.

### Calculation of F100 metric for *R. chloraquaticus*

*R. chloraquaticus* MAGs were used to calculate the F100 metric in a pairwise manner using the method developed recently (Conrad et al. 2024). Genomes for the Neutral Evolution Zero Recombination (NEZR) model were simulated using the code provided by Conrad and colleagues and were used for comparison against *R. chloraquaticus* and its lineages. Genomes were simulated in the ANI range 93-100% using a step size of 0.1 with 25 genomes at each of these values to construct the NEZR model. The *R. chloraquaticus* results, separated by genome quality and assigned lineage, were overlaid on the NEZR model to determine the importance of core gene content in *R. chloraquaticus* and the role of recombination in the maintenance of *R. chloraquaticus* lineages.

### Statistical analyses

All pairwise comparisons between groups for significance was calculated using the pairwise Wilcox test on R using the stats package.

## Results

### 3.1 Identification of good quality, contiguous genomes for R. chloraquaticus

Of the 42 unique MAGs representing *R. chloraquaticus*, 26 MAGs belonged to Lineage 1 and 16 MAGs to Lineage 2. These MAGs were analyzed either based on their individual (i.e., for pangenome analysis) or shared (i.e., for recombination analysis) gene content. The MAG quality criteria for both of these analyses are distinct and thus, a different number of MAGs were used for each analysis. The majority of the MAGs (41 out of 42) were more than 50% complete with less than 5% contamination (**Figure 1A)** and were all at least medium quality according to the MIMAG criteria (Bowers et al. 2017). However, this information is based on the presence/absence of marker genes and does not provide information on the quality and recovery of the remaining genes in the genome. This is important as several of the medium quality MAGs indicated high levels of fragmentation based on the number of contigs (**Figure 1B**), and the number of contigs was highly correlated with the number of partial/fragmented genes within the MAGs (**Figure 1C**). Fragmented assemblies can result in MAGs with higher number of partial genes, which can then affect the identification of shared (core) genes across MAGs and thus pangenomic and gene-based calculations. To account for this, we utilized the definition of high genome integrity from SeqCode, which identifies MAGs with less than 100 contigs as those with high genome integrity (Hedlund et al. 2022). Only 23 MAGs out of 42 had fewer than 100 contigs and a completeness greater than 80% and thus were defined as high genome integrity MAGs (**Figure 1B**). This filtering of MAGs mitigates issues of missing and partial genes that can hamper gene-based analysis especially for pangenomics.

**Figure 1:**
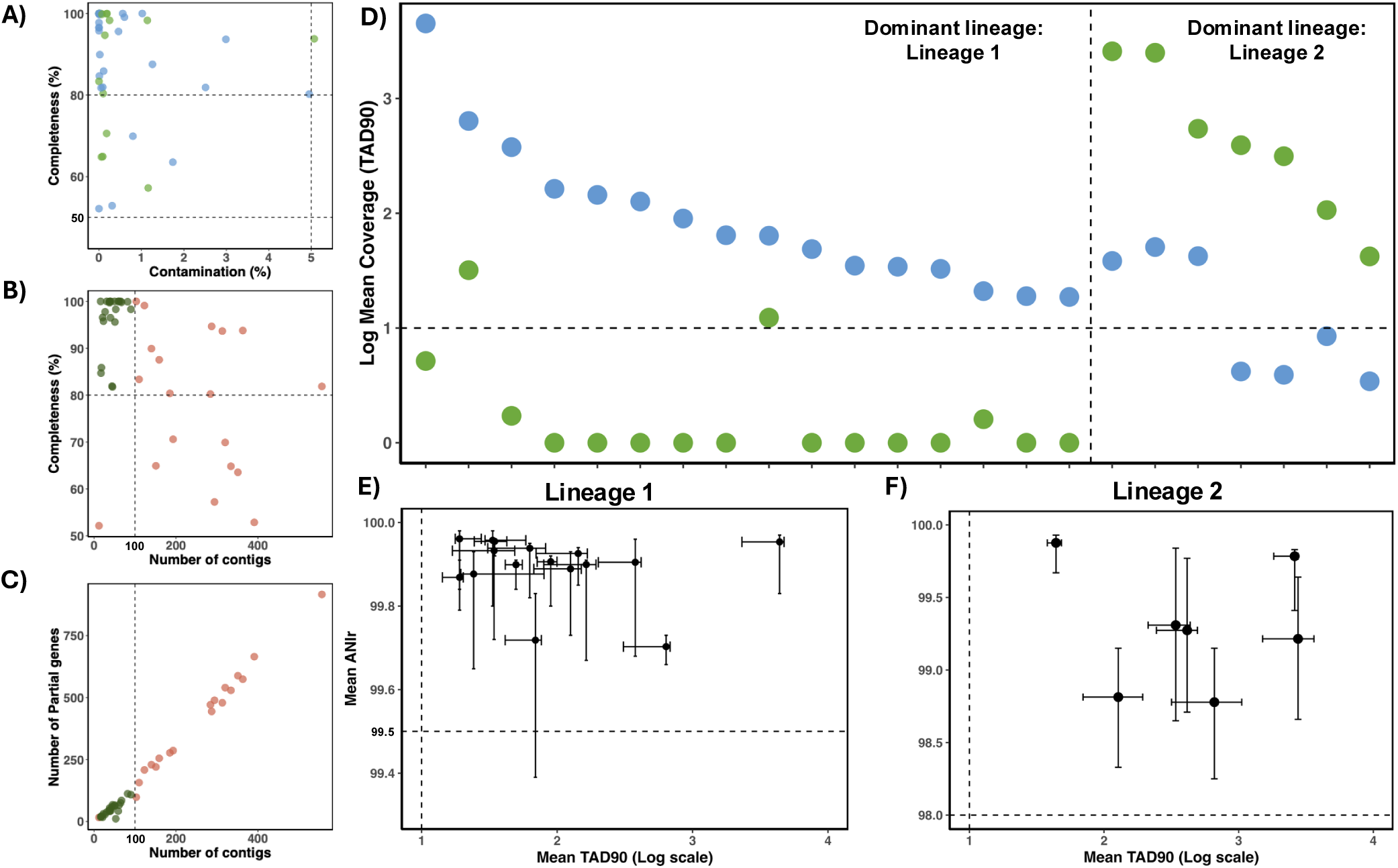
A comparison of (A) genome completeness and contamination, (B) completeness and number of contigs, and (C) number of partial genes and number of contigs of the 42 *R. chloraquaticus* MAGs. Colors represent the two lineages of *R. chloraquaticus*: Lineage 1 (Blue) and Lineage 2 (Green) in (A). Colors in (B) and (C) represent genome quality: low (Dark green) and high fragmented (Red) MAGs. D) Log mean coverage (TAD90) for the two *. R. chloraquaticus* lineages across the 23 DWDS from which high genome integrity MAGs were recovered. Colors represent the two lineages of *R. chloraquaticus* like Figure 1A. A comparison of mean ANIr and log mean TAD90 for DWDS, from where Lineage 1 (E) and Lineage 2 (F) MAGs were assembled.

An additional issue that could affect the conclusions of our MAG-based analysis is if the recovered *R. chloraquaticus* MAGs may be chimeric from the two lineages, which means it includes contigs representing both lineages within the same MAG. Such chimeric MAGs are much more likely to emerge from complex metagenomes if closely related organisms co-exist at similar relative abundances which limits the utility of differential abundance for binning purposes (Ramos-Barbero et al. 2019). To test for the possibility of chimeric MAGs, we mapped the reads from the 23 DWDS (from which the aforementioned high genome integrity MAGs were independently assembled) to two representative MAGs of the two lineages (**Figure 1D**). The two representative MAGs exhibited low fragmentation (31 contigs for Lineage 1 MAG and 59 contigs for Lineage 2 MAG), 100% completeness, and 0% contamination as determined by CheckM2. A significantly higher proportion of reads mapped to the MAG from the dominant lineage in each of these DWDS. In almost all cases (22 out of 23), the difference in sequencing depth coverage (or simply coverage) between the two lineages was at least an order of magnitude, suggesting that when these two lineages co-exist, differential abundance-based clustering should successfully bin their contigs in distinct MAGs. Furthermore, the coverage of the non-dominant lineage was consistently below 10x, the threshold necessary to obtain high quality MAGs with an expected breadth value of 100% (Lander and Waterman 1988), in 18 out of 23 DWDS (**Figure 1D**). In some DWDSs where both lineages were detected at substantial coverage (around 10x or more), we recovered two separate MAGs representing the two lineages. These results suggest that chimeric MAGs representing a hybrid of the two lineages is highly unlikely.

We further evaluated the extent to which the two MAGs (and their contigs) effectively represented the two lineages detected in DWDS. Previously, we observed that Lineage 1 consisted of organisms that are a part of the same genomovar (ANI>99.5%) and Lineage 2 consisted of a group of organisms that shared an ANI value of ∼98%. Therefore, we utilized two criteria to evaluate whether these lineages serve as good representatives of *R. chloraquaticus*: (1) all contigs of a MAG must have similar coverage (TAD90) and above the threshold of 10x where they can be effectively assembled and binned with an expected breadth value of 100% (Lander and Waterman 1988) in metagenomes that the group is detected, and (2) the ANIr values of contigs satisfy the lineage specific ANI thresholds of 99.5% ANI for Lineage 1 and 98% for Lineage 2. To do this, we mapped metagenomic reads from each DWDS to the representative MAG of the dominant lineage from the respective DWDS and estimated the TAD90 and ANIr to determine whether these lineages represent *R. chloraquaticus* well (**Figure 1E and 1F**). The reads mapping to contigs in the MAG representing the dominant lineage in each DWDS were consistently above 10x coverage indicating high confidence in the prevalence of these contigs with the average ANIr values within the ANI range for these lineages. This indicates that the two lineages appropriately represent the *R. chloraquaticus* organisms in these DWDS.

### 3.2 Lineage 1 and Lineage 2 organisms share a large core gene set while also showing lineage-specific gene signatures

We constructed a *R. chloraquaticus* pangenome with Lineage 1 and Lineage 2 serving as sub-classifications using the 23 high genome integrity MAGs to delineate core and accessory genes. Here, “core” genes represent those that are ideally present in all *R. chloraquaticus* organisms while “accessory” genes are only observed in a subset of these organisms and could consist of genes specific to either of the two lineages of *R. chloraquaticus* (Matthews et al. 2024). A major limitation of delineating “core gene set” based on their detection in genomes within a pangenome is that there are no established guidelines for gene detection threshold in order to account for uncertainty in detection. This problem is further exacerbated when MAGs are used to construct a pangenome where a missing gene could be due to lack of MAG completeness rather than being absent from a genome. To address this issue, we utilized a Bayesian approach to determine a core set of *R. chloraquaticus* genes which bypasses the need for prevalence cutoffs especially when using incomplete genomes and determines whether a gene is core or accessory based on likelihood ratios (Buck et al. 2022).

The *R. chloraquaticus* pangenome (Lineage 1: 16 MAGs and Lineage 2: 7 MAGs) was composed of 4272 gene clusters with 2628 clusters categorized as core genes and 1644 as accessory genes (**Figure 2A**). The MAGs used to construct the pangenome had an average size of 3.13 Mbp (range: 2.97-3.48 Mbp) and consisted of an average of 3028 genes (range: 2857-3351 genes). The core gene clusters make up a major portion of the pangenome (61.52%) and a major portion of the known gene content of any *R. chloraquaticus* MAG (Average value: 87.74%). In contrast, a large portion of the accessory gene clusters were only observed in less than three MAGs (51.58%) and a major portion of these are singleton gene clusters specific to individual MAGs(**Figure 2B**). The accessory gene clusters included 54 and 51 gene clusters that are unique to and only seen in all Lineage 1 and Lineage 2 MAGs, respectively.

**Figure 2:**
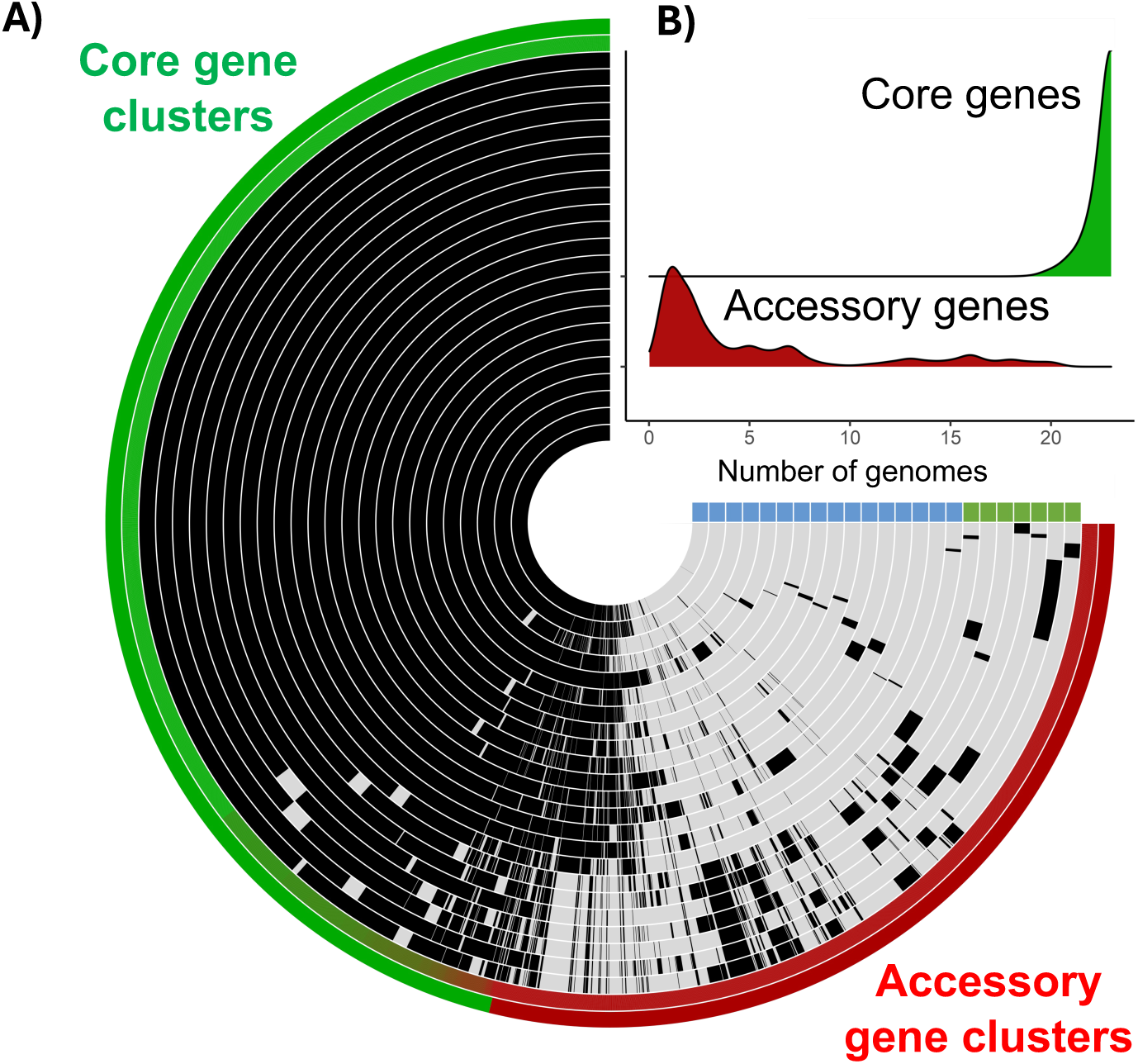
A) A schematic of the *R. chloraquaticus* pangenome. The concentric circles represent the 23 MAGs and are clustered according to their lineage assignment. B) Density plot depicting the prevalence of core (Green) and accessory (Red) gene clusters across the *R. chloraquaticus* MAGs.

Despite the difference in prevalence of the two *R. chloraquaticus* lineages, a large portion of their gene content is shared. This could indicate that the core set of genes described here are essential for the survival of *R. chloraquaticus* within disinfected DWDS. While the accessory gene cluster were mostly unique to specific MAGs, there were clusters that were unique to all MAGs of Lineage 1 and Lineage 2 and represented lineage specific gene clusters or signatures. These genes could likely explain the differential prevalence of the two lineages. In our previous study, we described several metabolic differences between the two lineages and how they can potentially contribute to difference in prevalence. For instance, the ability to oxidize thiosulfate was only detected in Lineage 1 but not in Lineage 2. Recycling sulfur compounds would be beneficial within an environment with reactive chlorine species stressors (Gray et al. 2013). Another example is the presence of the toxin-antitoxin system (HicAB) downstream of the chlorite dismutase gene in Lineage 1 and not in Lineage 2. This system is associated with persister/dormancy phenotypes and would allow cells to thrive in high stress conditions (Harms et al. 2018). Nonetheless, a large portion of the Lineage 1 specific gene clusters did not have a known function requiring further research into their role and if and how they contribute to fitness advantages.

### 3.3 *R. chloraquaticus* displays a relatively closed pangenome with Lineage 2 organisms displaying higher gene diversity than Lineage 1

Heap’s law describes a relationship between the number of gene clusters in a pangenome, and the number of genomes used to construct the pangenome. It is described by the following relationship:

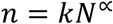

Where,

n = Number of gene clusters in the pangenome

N = Number of genomes used to construct the pangenome

α = Heap’s Law parameter

The Heap’s law parameter indicates whether a pangenome is open or closed. An open pangenome indicates that the addition of new genomes/MAGs to a pangenome analysis results in the addition of novel genes to the pangenome. In contrast, a closed pangenome indicates the number of novel genes added to a pangenome does not scale proportionally with the number of genomes and reaches a constant value as more genomes are sequenced. A Heap’s law parameter value less than 0 represents a closed pangenome and values greater than 0 would describe an open pangenome (Park et al. 2019). A Heap’s law parameter for all *R. chloraquaticus* MAGs (**Figure 3A**) of 0.12 indicates an open pangenome for this species. However, in comparison to other bacterial species identified in multiple ecosystems, *R. chloraquaticus* has a lower Heap’s law parameter value and a relatively “closed” pangenome (*E. coli*: 0.375, *M. tuberculosis*: 0.435, *P. aeruginosa*: 0.3491, *A. baumannii*: 0.3395) (Park et al. 2019). The value was low even when compared to organisms that are adapted to very select ecosystems (SAR11: 0.34, *S. pepae*: 0.44) (Grote et al. 2012, Viver et al. 2023). Overall, these results and comparisons suggest that *R. chloraquaticus* has a highly conserved set of genes that likely aid in its adaptation in disinfected DWDS.

**Figure 3:**
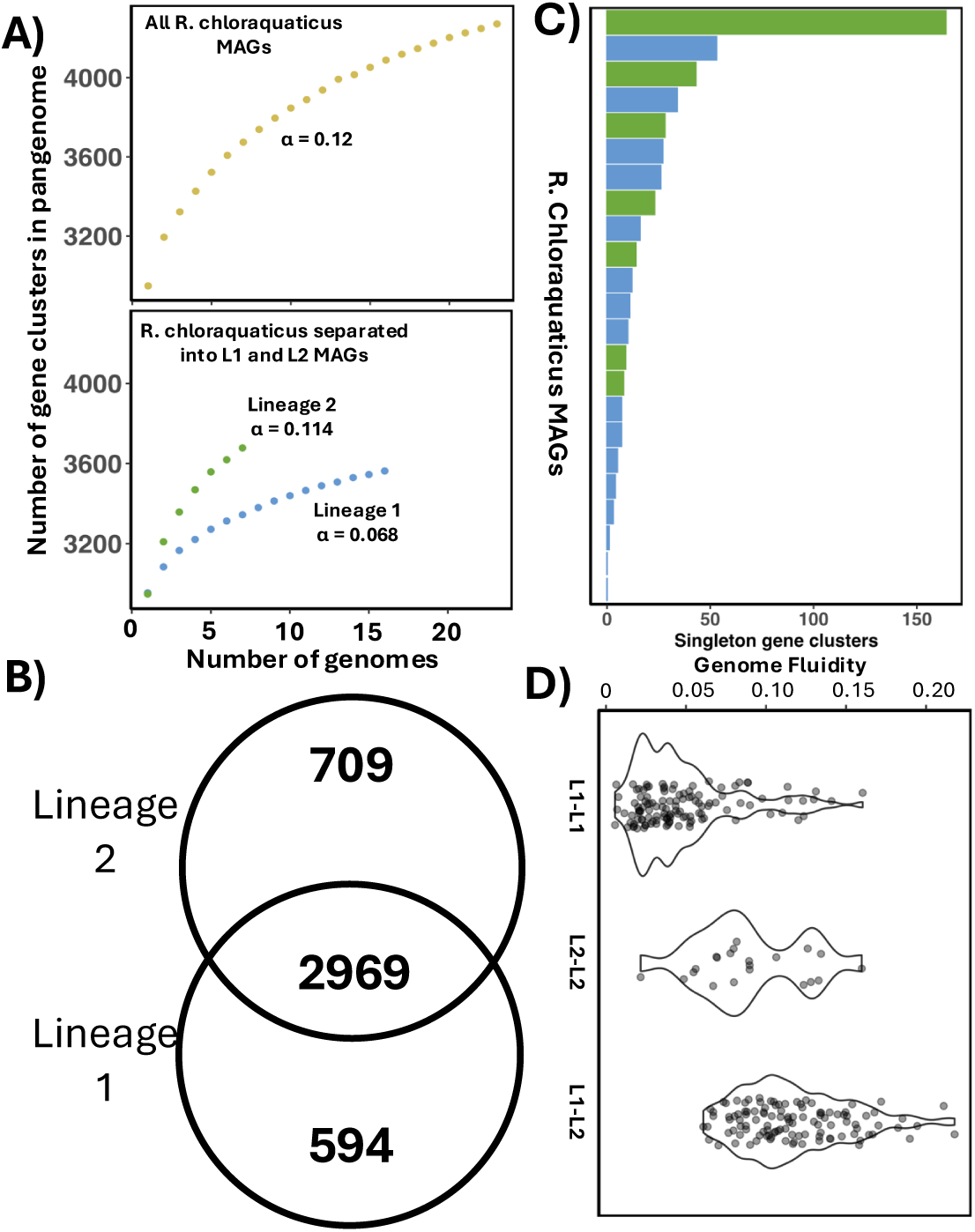
A) The pangenome structure of the *R. chloraquaticus* lineages. B) A Venn diagram depicting the distribution of gene clusters between the two lineages. C) Number of singleton gene clusters observed in the *R. chlora*quaticus MAGs used to in the pangenome analysis. D) Genome fluidity for intra- and inter lineage comparisons within *R. chloraquaticus*.

Focusing the pangenome analysis at the lineage-level indicates that Lineage 2 has a higher gene diversity (Heap’s law parameter: 0.114) as compared to Lineage 1 (Heap’s law parameter: 0.069). These results also indicate that most of the gene diversity within *R. chloraquaticus* pangenome is due to Lineage 2 organisms despite the fact that more Lineage 1 MAGs were used in this analysis (16 Lineage 1 MAGs Vs 7 Lineage 2 MAGs) (**Figure 3B**). These results are also consistent with the fact that Lineage 2 harbors more genomic diversity in terms of number of unique genomovars and/or lower ANI values, and the relationship between (greater) gene content differences with (lower) ANI values observed even at the intraspecies level (Rodriguez-R et al. 2023). Analysis of singleton gene clusters indicates that a few Lineage 2 MAGs harbored disproportionately more singleton gene clusters such as one Lineage 2 MAG (MAG ID: RCL_L2_12) harbored 164 singleton genes (**Figure 3C**).

Determining the core gene content of a taxonomic group using pangenomics is limited by the number of genomes/MAGs used to construct a pangenome. Additionally, the core gene content analysis and Heap’s law analysis cannot be used to quantify the differences in gene content between organisms of the same taxonomic group. To address this issue and quantify the differences, we utilize a metric called genome fluidity (Kislyuk et al. 2011). It is described by the equation shown below:

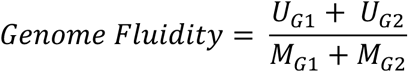

Where,

U_G1_ = Gene clusters unique to MAG 1 (G1)

U_G2_ = Gene clusters unique to MAG 2 (G2)

M_G1_ = Total gene clusters observed in MAG 1 (G1)

M_G2_ = Total gene clusters observed in MAG 2 (G2)

This metric is not dependent on the number of genomes used for the analysis and is used to determine differences in gene content between two organisms within a taxonomic group. This allows quantitative determination of the percentage of the known gene content would be unique between two genomes of *R. chloraquaticus* and its lineages. Mean genome fluidity estimates from pairwise comparisons of MAGs within Lineage 1 and Lineage 2 were 4.77% and 8.97%, respectively (**Figure 2D**). These results indicate that these two lineages clearly show a difference in terms of gene diversity with lower gene diversity in Lineage 1 relative to Lineage 2. These genome fluidity results also indicate that the retrieval of additional genomes from Lineage 1 organisms would only yield a limited number of new/novel genes and that a major portion of the gene families of Lineage 1 organisms has been captured by the MAGs included in this study.

Overall, despite a relatively closed pangenome and a low gene diversity, *R. chloraquaticus* lineages and genomovars are globally detected within disinfected DWDS suggesting a form of selection. One possible reason could be genome streamlining. Genome streamlining involves trade-off where organisms loose unnecessary gene content and functions in order to thrive better within one ecosystem (Giovannoni et al. 2014). Two lines of evidence that support this statement are: (a) low gene diversity as seen through the relatively closed pangenome and low genomic fluidity and (b) *R. chloraquaticus* MAGs have only been obtained and observed in disinfected drinking water systems thus far. It should be mentioned, however, that MAGs are known to underrepresent the accessory genes of the genome due to technical limitations during the assembly and/or binning processes (Meziti et al. 2021). Accordingly, we focused our analysis on high-quality and high genome integrity MAGs only, and we believe that our conclusions about the relative gene fluidity or pangenome sizes among the two lineages and related findings are robust. Nonetheless, it is likely that our estimate of how large the core gene set is as a proportion of the total genome represents an overestimate due to the above-mentioned limitation.

### 3.4 Homologous recombination and geographic isolation are potential mechanisms for the maintenance of genome diversity within *R. chloraquaticus*

Beyond gene diversity, gene-level similarity analysis is critical for inferring the strength of environmental selection pressures, while also revealing the extent of homologous recombination between closely related organisms. To this end, we utilized the F100 metric to estimate the number of identical genes expected between any two genomes based on their ANIs under a neutral evolution zero recombination (NEZR) model and compare it to the observed number of such identical genes. The difference between the two numbers serves as a proxy for the frequency or magnitude of recent homologous recombination, as proposed recently (Conrad et al., 2024). We determined the F100 values for *R. chloraquaticus* MAGs to (a) evaluate the extent to which the core gene set is identical within different lineages and (b) determine the extent of homologous recombination using the F100 as a proxy metric and in comparison, with the NEZR model. Both aspects are critical for contextualizing the role of the core gene content in *R. chloraquaticus* and its selective detection in disinfected DWDS globally.

The F100 values for pairwise comparisons of Lineage 1 MAGs were significantly higher (P<2*10^-16^, pairwise Wilcox test) than the NEZR model (**Figure 3A**) irrespective of the quality of the MAGs being compared. The F100 values estimated using the NEZR model for genomes within the same genomovar (ANI:99.53-99.91%) ranged between 22.08-62.28% (mean: 39.38%) identical shared genes between MAGs being compared. In contrast, the proportion of identical genes for *R. chloraquaticus* Lineage 1 MAGs was 52.23-98.89% (mean: 88.4%). This significantly higher proportion of shared identical genes was consistently observed irrespective of the quality of the MAGs being compared. While homologous recombination could potentially explain the large proportion of identical genes, it is difficult to arrive at this conclusion at high levels of genome similarity as is seen in *R. chloraquaticus* Lineage 1 MAGs (ANI: 99.498 – 99.995%). However, the detection of a large proportion of identical core genes further reaffirms that these could be potentially associated with adaptation to disinfected drinking water systems and explain the global prevalence of Lineage 1 organisms. Intra Lineage 2 comparisons result in more variable F100 values (F100 Range: 1.75-93.79%) as compared to Lineage 1 (F100 range: 52.23-98.89%). This variability is expected considering Lineage 2 represents a more diverse group of organisms with shared ANI of ∼98% as opposed to Lineage 1 (ANI>99.5%). The F100 for Lineage 2 were not significantly different (P=0.428, Pairwise Wilcox Test) from those obtained from an NEZR model derived from a similar ANI range (ANI: 97.23-99.91%) when comparing less fragmented MAGs against each other. A possible explanation could be that MAGs from different genomovars within Lineage 2 could have emerged through random mutations. Lineage 2 consists of four separate clusters with each with an ANI range of 99-99.91% and this could likely constitute four different genomovars. Within genomovar comparisons for Lineage 2 yielded significantly higher F100 values (P< 2*10^-16^ Pairwise Wilcox Test) as compared to NEZR model, while cross genomovar comparisons resulted in F100 values that were not significantly different from the NEZR model (P=0.4 Pairwise Wilcox Test) (**Figure 4B**). Despite these observations, multiple individual comparisons within Lineage 2 had F100 values higher than the expected F100 values determined through the NEZR model for a given ANI value (**Figure 4A**). This indicates that recombination likely occurs in specific instances even if it is unlikely to be a universal feature of the group. However, the high degree of fragmentation and higher diversity within Lineage 2 MAGs hinders the ability to draw conclusions based on a subset of these values. Further experimentation can determine the prevalence of recombination within Lineage 2 organisms.

**Figure 4:**
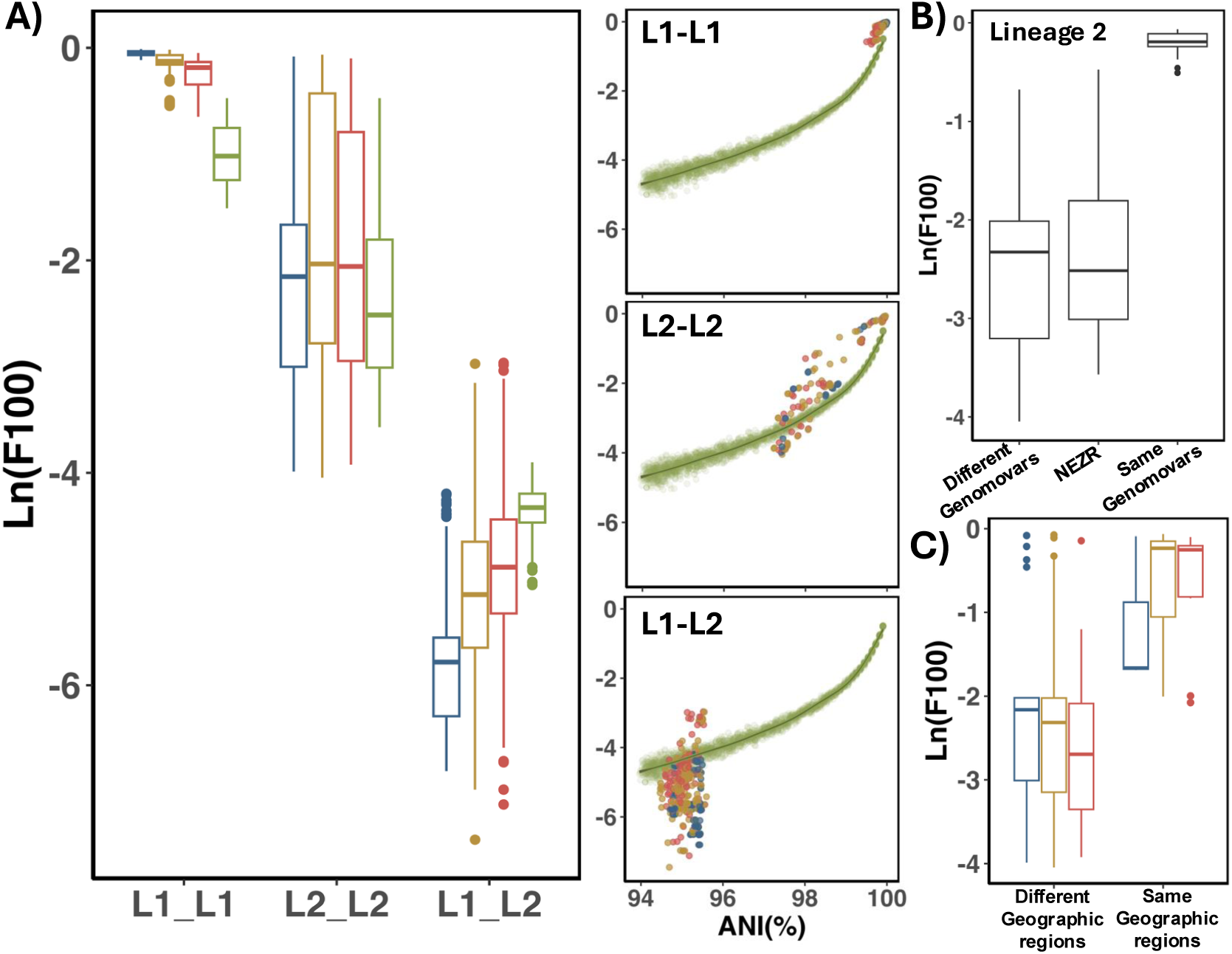
A) Log F100 boxplot for inter and intra lineage comparisons of *R. chloraquaticus*. Boxplot is split according to quality: high genome integrity – high genome integrity MAG comparisons (blue), high genome integrity – low genome integrity MAG comparisons (yellow) and low genome integrity – low genome integrity MAG comparisons (red) and NEZR values (green). Ln(F100) variance with ANI depicts the inter and intra lineage comparisons of *R. chloraquaticus* and the NEZR model. Colors depict the comparisons by MAG quality. B) Ln(F100) comparisons within R. chloraquaticus (Lineage 2) separated according to inter and intra-genomovar comparisons. C) Ln(F100) comparisons within *R. chloraquaticus* Lineage 2 MAGs separated according to geographic region and quality of MAGs. Colors depict MAG quality used for comparison similar to Figure 4A.

While Lineage 1 is a globally prevalent genomovar, Lineage 2 consisted of organisms seen both globally as well as ones that were restricted to certain geographic regions. Overall, F100 values estimated for Lineage 2 MAGs from the same geographic region were significantly higher (P = 2*10^-11^, Pairwise Wilcox Test) than those estimated when comparing Lineage 2 MAGs from different regions (**Figure 4C**). Therefore, it is likely that geographic isolation could also plays a role in creating diversity within *R. chloraquaticus*. Comparing MAGs from Lineage 1 and Lineage 2 resulted in F100 values that were significantly lower than the NEZR model (P<2*10^-16^ Pairwise Wilcox Test). These results would indicate that it is unlikely that homologous recombination occurs between the two lineages of *R. chloraquaticus*. This is also consistent with the observation that the two lineages of *R. chloraquaticus* did not co-occur together in most DWDS included in this study. Based on the F100 results, ecological separation and the whole-genome similarity between the two lineages (∼95% ANI), it is highly probable that these two lineages represent independent, but closely related species. However, due to the environmental relevance of the two lineages and the lack of phenotypic information about these groups, we retain our categorization of these two lineages as sub-species within *R. chloraquaticus* until further studies elucidate phenotypic and ecological differences between them.

### Conclusions

Through systematic pangenome analysis we demonstrate that *R. chloraquaticus* possesses a core set of genes that constitutes a majority of the known gene content of the genome. Further, these core genes demonstrate high levels of sequence homology in Lineage 1, the globally dominant genomovar, compared with Lineage 2. Lineage 2 also exhibits higher gene diversity relative to Lineage 1 and comprises of geographically restricted genomovars. While there are functionally relevant differences between the two lineages, our combined analyses suggest potential adaptation of *R. chloraquaticus* to an ecosystem as unique as the disinfected DWDS that is accompanied by genome streamlining, especially in Lineage 1. Considering the previously reported high level of relative abundance and detection frequency of *R. chloraquaticus* in disinfected DWDS and gene-level adaptation of *R. chloraquaticus* Lineage 1 reported in this study, organisms from *R. chloraquaticus* Lineage 1 could represent model organisms for future efforts to elucidate how drinking water treatment and management-induced selective pressures shape the drinking water microbiome.

## Acknowledgement

This research was supported by the National Science Foundation (NSF) (CBET Award Number: 2220792) and US Environmental Protection Agency (Grant R840606).

## Data availability

The non-redundant MAGs that form the basis of the DWGC and species-level representative MAGs are available on FigShare (DOI: https://doi.org/10.6084/m9.figshare.c.7245403.v1).

